# Kaposi’s Sarcoma-associated Herpesvirus Promotes Mesenchymal-to-Endothelial Transition by Resolving the Bivalent Chromatin of PROX1 Gene

**DOI:** 10.1101/2021.04.24.441268

**Authors:** Yao Ding, Weikang Chen, Zhengzhou Lu, Yan Wang, Yan Yuan

## Abstract

Human mesenchymal stem cells (MSCs) are highly susceptible to Kaposi’s sarcoma-associated herpesvirus (KSHV) infection, and the infection promotes mesenchymal-to-endothelial transition (MEndT) and acquisition of Kaposi’s sarcoma (KS)-like phenotypes. Increasing evidence suggests that KS may arise from KSHV-infected MSCs. To understand how KSHV induces MEndT and transforms MSCs to KS cells, we investigated the mechanism underlying KSHV-mediated MSC endothelial lineage differentiation. Like embryonic stem cells, MSC differentiation and fate determination are under epigenetic control. Prospero homeobox 1 (PROX1) is a master regulator that controls lymphatic vessel development and endothelial differentiation. We found that the PROX1 gene in MSCs harbors a distinctive bivalent epigenetic signature consisting of both active marker H3K4me3 and repressive marker H3K27me3, which poises expression of the genes, allowing timely activation upon differentiation signals or environmental stimuli. KSHV infection effectively resolved the bivalent chromatin by decreased H3K27me3 and increased H3K4me3 to activate the PROX1 gene. vIL-6 signaling leads to the recruitment of MLL2 and Set1 complexes to the PROX1 promoter to increase H3K4me3, and the vGPCR-VEGF-A axis is responsible for removing PRC2 from the promoter to reduce H3K27me3. Therefore, through a dual signaling process, KSHV activates PROX1 gene expression and initiates MEndT, which renders MSC tumorigenic features including angiogenesis, invasion and migration.

## Introduction

Kaposi’s sarcoma-associated herpesvirus (KSHV), also termed human herpesvirus type 8 (HHV8), is a member of the γ-herpesviridae subfamily (Chang et al., 1994). This virus has been proven to be the etiological agent of Kaposi’s sarcoma (KS) (Moore and Chang, 1995), primary effusion lymphoma (PEL) (Cesarman et al., 1995), and multicentric Castleman’s disease (MCD) (Soulier et al., 1995). Recently, KSHV was found to be associated with childhood osteosarcoma (Chen et al., 2021). However, the lack of comprehensive understanding about KSHV infection and tumorigenesis hampers our ability to treat KSHV-associated cancers. For example, KS is atypical cancer. The onset of KS as simultaneously multiple lesions (in the absence of obvious metastasis), the multiclonal malignant nature (especially in the late stages), and the presence of unusual inflammatory cell infiltration and neoangiogenesis in the very early stage of KS distinguish KS from other orthodox tumors. The origin of KS spindle cells remains elusive. The current leading model is that KS spindle cells may derive from endothelial cell lineage as KS cells express endothelial markers. However, KS cells are poorly differentiated and also express other markers such as smooth muscle, macrophage, and mesenchymal markers, suggesting that KS cells do not faithfully represent the endothelial cell lineage (Cancian et al., 2013). Recently, we found a series of evidence suggesting that KS originates from oral mesenchymal stem cells (MSCs) through KSHV-induced mesenchymal-to-endothelial transition (MEndT) (Li et al., 2018).

MSCs have been identified as a population of hierarchical postnatal stem cells with the potential to self-renew and differentiate into osteoblasts, chondrocytes, adipocytes, cardiomyocytes, myoblasts, and neural cells (Friedenstein et al., 1974; Prockop, 1997). The oral cavity contains a variety of distinct MSC populations, including dental pulp stem cells (DPSCs), periodontal ligament stem cells (PDLSCs), and gingiva/mucosa-derived mesenchymal stem cells (GMSCs) (Gronthos et al., 2000; Miura et al., 2003; Zhang et al., 2009). Among these MSCs, PDLSCs and GMSCs have the potential to directly interact with oral cavity saliva, microbiota, and virus, therefore having a great chance to be infected by KSHV in the oral cavity. Our recent study found that AIDS-KS spindle cells express <u>Ne</u>uroectodermal <u>st</u>em cell marker (Nestin) and oral MSC marker CD29, suggesting an oral/craniofacial MSC lineage of AIDS-associated KS (Li et al., 2018). Furthermore, MSCs were found highly susceptible to KSHV infection, and the infection effectively promotes multiple lineage differentiation, especially endothelial differentiation. When implanted in mice, KSHV-infected MSCs were transformed into KS-like spindle-shaped cells with other KS-like phenotypes (Wang et al., 2020). These findings provided evidence that KS derives from KSHV-infected MSCs. In addition, other laboratory identified PDGFRA+ MSCs as KS progenitors (Naipauer et al., 2019).

We sought to understand how KSHV infection regulates the MEndT process to promote malignant transformation. It is known that the regulation of stem cell fate determination and differentiation is mainly at the epigenetic level, which has been intensively studied in embryonic stem cells (Christophersen and Helin, 2010). Embryonic stem cells undergo global remodeling during early stem cell development that commits them to the desired lineage. The lineage commitment of stem cells is controlled by epigenetic mechanisms, and histone modification appears to be the most important layer in such regulation (Kouzarides, 2007). Some critical transcription factor gene promoters in embryonic stem (ES) cells harbor a distinctive histone modification signature that combines the activating histone H3K4me3 mark and the repressive H3K27me3 mark. These bivalent domains are considered to poise the expression of developmental genes, allowing rapid activation upon developmental signals (Bernstein et al., 2006; Voigt et al., 2013). Increasing evidence suggests that mesenchymal stem cell differentiation, like embryonic development regulation, is also controlled at the epigenetic level (Pérez-Campo and Riancho, 2015). Since MSCs face the same challenge as embryonic stem cells –– multi-lineage differentiation and stem cell fate determination, it is possible that similar epigenetic mechanism found in embryonic stem cells may also apply to MSCs. The homeobox gene Prospero homeobox 1 (PROX1) is a master regulator gene that controls lymphatic vessel development and endothelial differentiation (Wigle et al., 2002) and is up-regulated by KSHV infection in MSCs (Li et al., 2018). Elucidation of the regulation mechanism of PROX1 gene expression by KSHV is vital for comprehension of KSHV-mediated MEndT that leads to KS.

In the current study, we found that the PROX1 gene in MSCs exhibits a “bivalent” epigenetic signature consisting of both H3K4me3 and H3K27me3 marks. We investigated how the bivalent chromatin structure is established and how KSHV infection resolves the bivalent domain to activate PROX1 and MEndT genes. This study elucidates the mechanism underlying KSHV-mediated MEndT at the transcription level.

## Results

### Kaposi’s sarcoma lesions express PROX1 and KSHV infection of mesenchymal stem cells induces PROX1 expression

It was reported that among 30 oral Kaposi sarcomas (KS) lesions, twenty-eight (93.3%) were positive for PROX1 (Lavado and Oliver, 2007). To confirm that KSHV infection is responsible for the induction of PROX1 expression, we examined three cases of KS, one oral and two skin lesions for expression of PROX1 and KSHV antigen LANA using immunohistochemistry analysis. The result showed that all three samples were PROX1-positive and LANA positive (Fig. 1A). Periodontal ligament stem cells (PDLSCs) were infected with KSHV and the expression of PROX1 in response to KSHV was examined by double-staining immunofluorescence assay (IFA) and Western analysis. IFA showed that PROX1 expression was consistently correlated with KSHV infection (LANA expression) (Fig. 1B), and Western blot analysis showed that PROX1 expression was upregulated by KSHV (Fig. 1C). Furthermore, mock- and KSHV-infected PDLSCs were implanted in immunocompromised mice under the kidney capsule. After four weeks, the implants were subjected to an immunohistochemistry study for PROX1 expression. As shown in Fig. 1D, PROX1 expression was found to be significantly elevated in KSHV-infected PLDSCs in comparison to mock-infected PDLSCs. Overall, PROX1 expression is upregulated both *in vivo* and *in vitro* in response to KSHV infection in MSCs.

**Fig 1.**
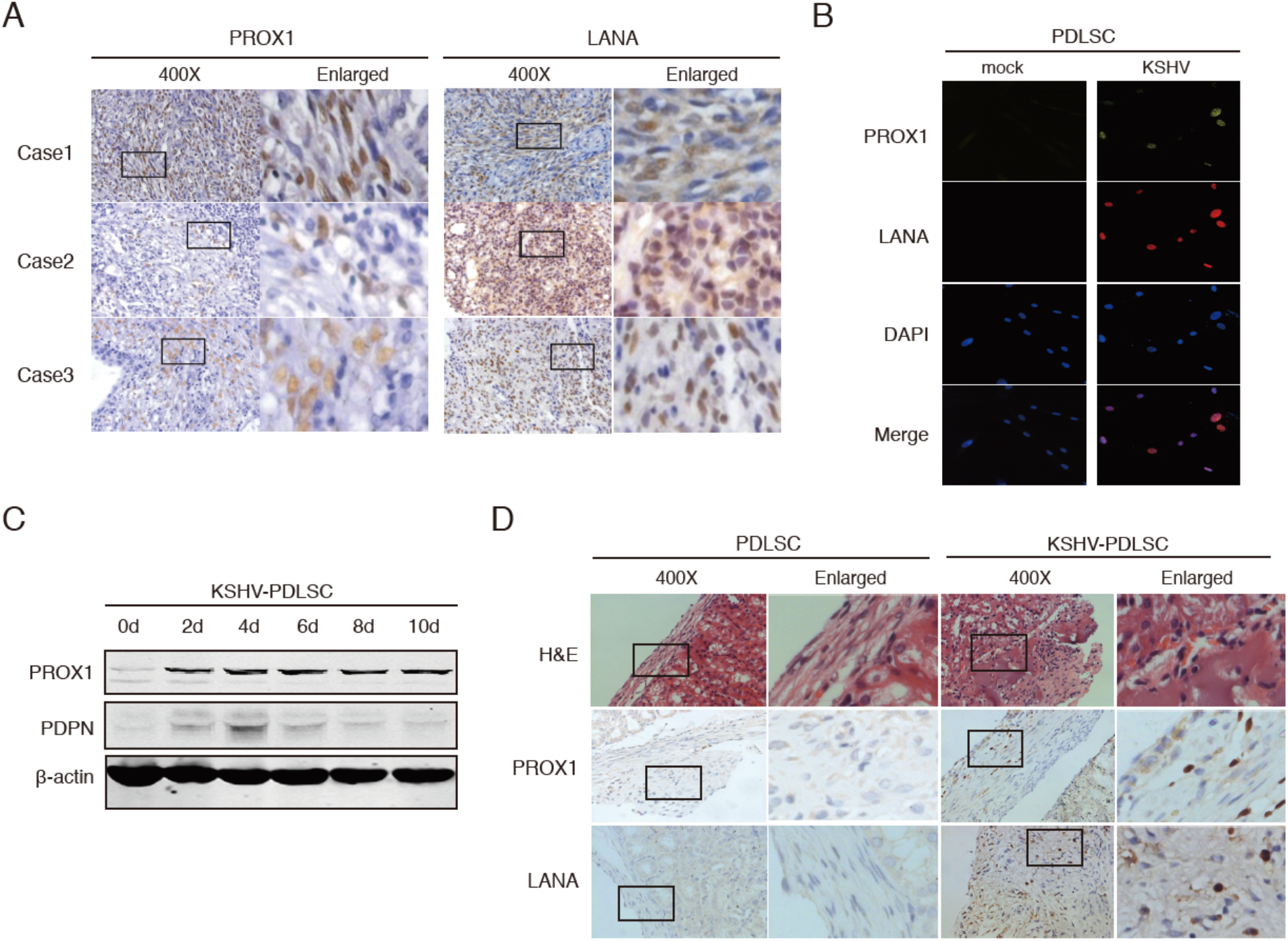
Kaposi’s sarcoma lesions express PROX1 and KSHV infection of mesenchymal stem cells induces PROX1 expression. (A) Paraffin-embedded sections of Kaposi sarcoma (KS) lesions from three AIDS-KS patients were analyzed by immunohistochemistry (IHC) staining (400x and enlarged) for PROX1 and LANA. (B) Mock- and KSHV-infected PDLSCs were examined the expression of PROX1 and LANA by immunofluorescence assay (IFA). (C) Time courses of the expression of PROX1 and PDPN in PDLSCs after KSHV infection. (D) mock- and KSHV-infected PDLSCs were transplanted in mice under kidney capsules. After 4 weeks, PROX1 and LANA expression in the implants was analyzed by IHC.

### PROX1 expression is essential for KSHV-induced mesenchymal-to-endothelial transition

PROX1 is the master regulator of endothelial differentiation and lymphatic vessel development. The expression of PROX1 in KS lesions and the induction of PROX1 expression by KSHV in PDLSCs suggest that PROX1 plays a critical role in KSHV-mediated MEndT and KS development. To explore this hypothesis, we determined the role of PROX1 in MEndT by using an shRNA-mediated silencing approach. An shRNA specific to PROX1 was introduced into PDLSCs by a lentiviral vector, and then the transduced PDLSCs were infected by KSHV. The knockdown efficiency of the shRNA was validated by Western blot (Fig. 2A). The contribution of PROX1 to endothelial lineage differentiation and neovascularity was evaluated by Western blot to test endothelial markers expression and a Matrigel tubulogenesis assay. Results showed that the expression level of lymphatic endothelial markers (VEGFR3, LYVE-1, and PDPN) and pan-endothelial marker (VEGFR2) decreased in PDLSCs transduced with shPROX1 compared to those with control shRNA (Fig. 2B). Correspondingly, shRNA-mediated PROX1 inhibition resulted in the loss of tubulogenesis (Fig. 2C). Transwell assay was used to assess cell invasion driven by PROX1. The result showed that shRNA-mediated PROX1 inhibition significantly reduced invasion ability compared to control shRNA (Fig. 2D). Furthermore, when implanted in mice under kidney capsules, KSHV-infected PDLSCs underwent an MEndT process revealed by the expression of endothelial markers CD31, VCAM1, and PDPN, while MEndT did not occur in PROX1-knockdown cells as they failed to express those endothelial markers (Fig. 2E), suggesting that PROX1 is essential for the initiation of MEndT induced by KSHV.

**Fig. 2.**
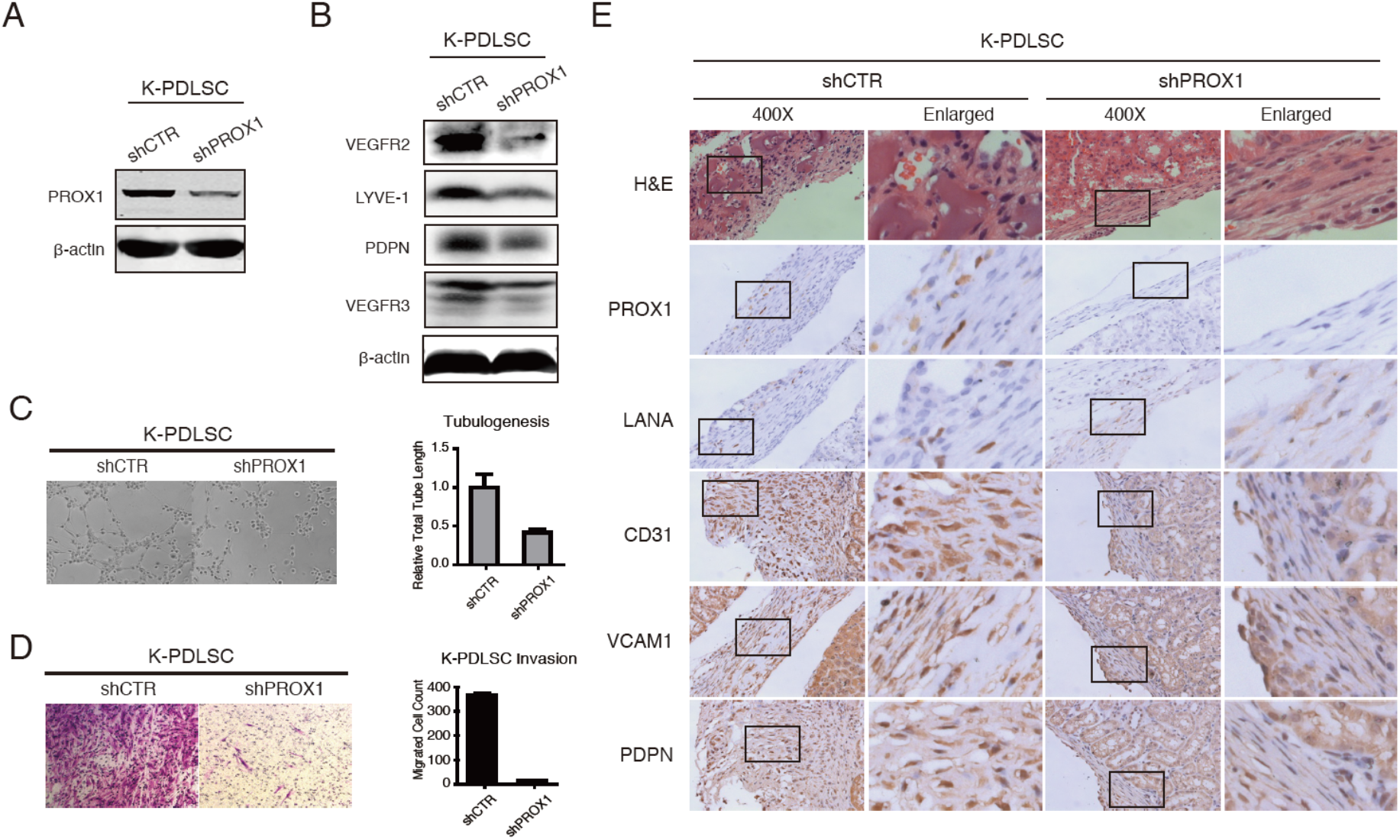
The critical role of PROX1 in mesenchymal-endothelial transition induced by KSHV. An shRNA lentivirus specific to PROX1 (shPROX1), along with a control shRNA (shCTR), were transduced into PDLSCs, followed by infection with KSHV. (A) The efficiency of PROX1 silencing was verified by Western blot. (B) The effects of the shRNA on the expression of pan-endothelial and lymphatic endothelial markers VEGFR2, LYVE-1, PDPN, and VEGFR3 were examined by Western blot. (C) KSHV-infected PDLSCs (K-PDLSC) transduced with shPROX1 or shCTR were evaluated for endothelial differentiation and angiogenic activities through a tubule formation assay. (D) K-PDLSCs transduced with shPROX1 or shCTR were subjected to cell invasion assay to determine the effect of PROX1 silencing on cell migration and invasion ability of K-PDLSCs. (E) KSHV-PDLSCs transduced with shPROX1 or shCTR were implanted under the kidney capsule of immunocompromised mice for 28 d. Then the kidney capsule was taken out and analyzed by hematoxylin and eosin (H&E) and IHC (PROX1, LANA, CD31, VCAM1, and PDPN) staining.

### KSHV promotes MEndT through epigenetic regulation of the PROX1 gene that possesses a bivalent chromatin structure

We sought to understand how KSHV infection regulates the MEndT process. Evidence suggests that mesenchymal stem cell differentiation is controlled at the epigenetic level (Pérez-Campo and Riancho, 2015). KSHV-mediated MSCs reprogramming likely takes place at this level. Histone modifications play critical roles in regulating the expression of developmental genes in mammalian development (Li, 2002; Bogliotti and Ross, 2012). To explore epigenetic regulation of KSHV-mediated MEndT, we examined histone modifications on the chromatin of several MEndT genes in mock- and KSHV-infected PDLSCs using Chromatin immunoprecipitation (ChIP) assay. Results revealed an interesting feature that PROX1 gene locus in mock-infected PDLSCs heavily consists of both repressive mark H3K27me3 and active mark H3K4me3. KSHV-infection dramatically increased H3K4me3 and decreased H3K27me3 (Fig. 3A). This chromatin structure pattern, known as bivalent chromatin structure, has been reported in the genes of key transcriptional factors responsible for development in embryonic stem cells (Bernstein et al., 2006). In contrast, other MEndT relevant genes, such as PML, TGF-β3, and TGF-βRII, obtained elevated H3K4me3 in the chromatin of their regulatory regions in response to KSHV infection (Fig. 3A).

**Fig. 3.**
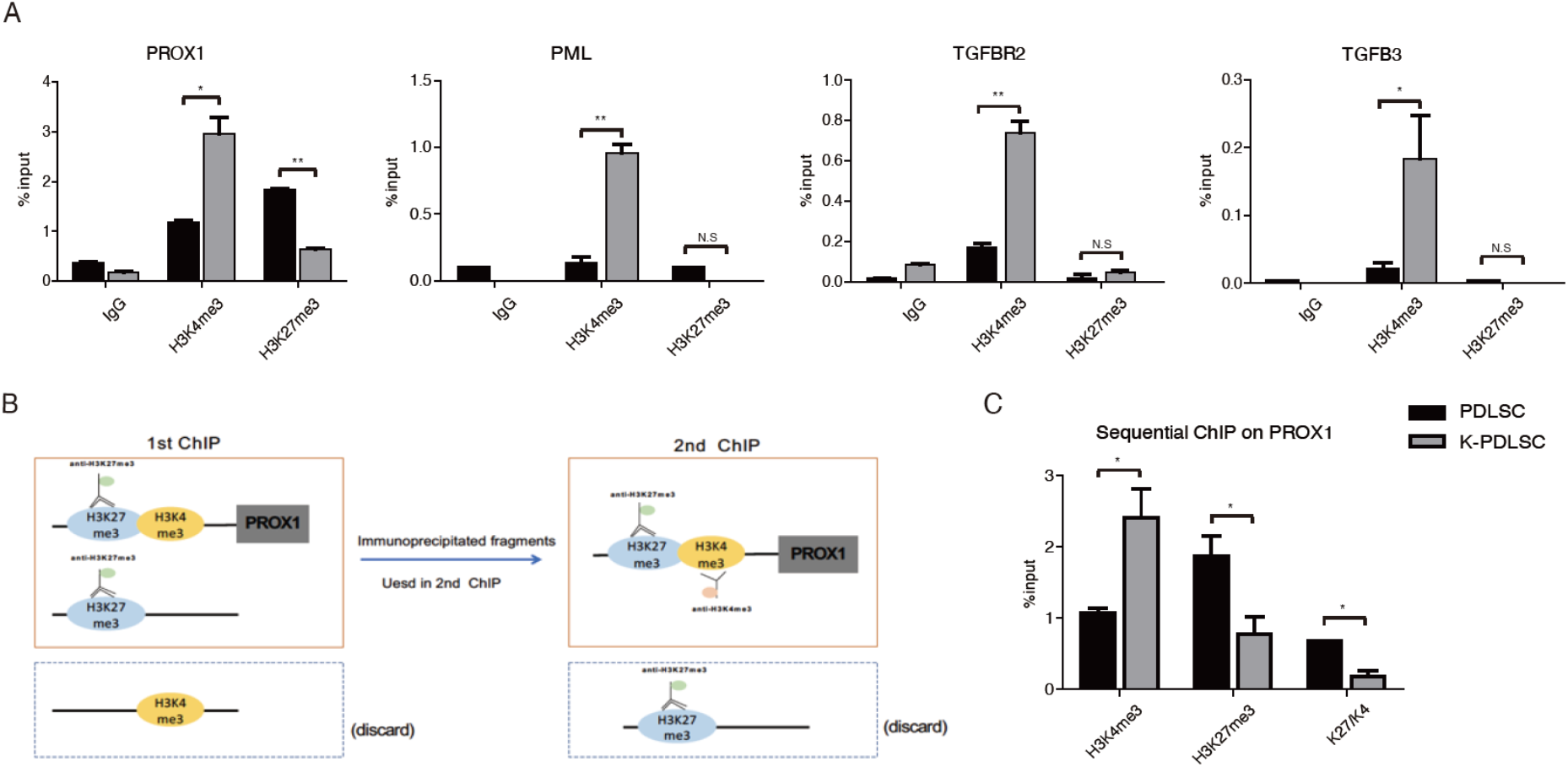
PROX1 gene is harbored in a bivalent chromatin structure and KSHV infection can resolve it to activate PROX1 expression. (A) Histone modifications (H3K4me3 and H3K27me3) of MEndT-related genes in mock- and KSHV-infected PDLSCs were determined by ChIP assay. PROX1 gene in mock-infected PDLSCs heavily consists of both active mark H3K4me3 and repressive mark H3K27me3, exhibiting a distinctive bivalent chromatin domain. (B) Schematic illustration of the sequential ChIP assay. Cross-linked chromatin from MSCs was immunoprecipitated with an anti-H3K27me3 antibody followed by a second immunoprecipitation with an anti-H3K4me3 antibody. (C) The extracted DNA was analyzed by RT-qPCR with primers of the PROX1 promoter, and the result showed that H3K4me3 and H3K27me3 co-exist in the same nucleosome in the PROX1 promoter. Error bars represent SD. Statistics analysis was performed using one-way ANOVA test, and P-value was calculated by GraphPad Prism. P-value < 0.05 was considered significant (*P<0.05, **P<0.01 and ***P<0.001) and N.S represented no significance.

Given the unique nature of the bivalent domain on the PROX1 gene, we would confirm that the observed bivalent structure truly reflects the simultaneous presence of both H3K4me3 and H3K27me3 on the same chromosome rather than the presence of two subpopulations with different epigenetic characters. To this end, we performed a sequential ChIP assay in which PDLSCs chromatin was immunoprecipitated first with the Lys27 tri-methyl antibody and secondly with the Lys4 tri-methyl antibody (Fig. 3B). This sequential purification is designed to retain only chromatin that concomitantly carries both the active and repressive histone modifications. Then real-time PCR was used to detect PROX1 promoter DNA on the bivalent nucleosomes. As shown in Fig. 3C, H3K4me3 and H3K27me3 were found to co-exist on the same PROX1 promoter chromatin, accurately representing a true bivalent epigenetic state. Overall, our result suggests that the bivalent epigenetic feature may silence the PROX1 gene and its downstream cascade in MSCs but keep them poised for activation. KSHV infection disrupts the balance and MSCs quickly turn on the PROX1 gene and downstream cascade, leading to MEndT.

### KSHV resolves the PROX1 bivalent chromatin by recruiting MLL2 and Set1 complexes to and removing PRC2 from the PROX1 promoter

To understand how KSHV manipulates PROX1 epigenetic regulation, we aimed to investigate the mechanism for establishing the bivalent domain in the PROX1 promoter and resolving it by KSHV. The central players in setting up and maintaining bivalent chromatin structure are the trithorax group (TrxG) and polycomb group (PcG) proteins (Voigt et al., 2013). The former consists of SET1A/B and MLL1-4 complexes, catalyzing the trimethylation of histone H3 Lys4 (Shilatifard, 2012). The latter forms multi-subunit polycomb-repressor complexes (PRC) 1 and 2, responsible for the trimethylation of histone H3 Lys27. PRC2 is particularly crucial for H3K27me3 in many developmental genes. To identify the tri-methyltransferase complexes that are recruited to PROX1 chromatin and altered in response to KSHV infection, ChIP assays were performed on mock- and KSHV-infected PDLSCs with antibodies against CFP1, MLL1, 2, respectively. The results showed that (i) prior to KSHV infection, MLL1 had been recruited to the PROX1 promoter region; (ii) KSHV infection did not change MLL1 content in the chromatin but recruited MLL2 and CFP1 to the promoter (Fig. 4A-C). CFP1, a CXXC finger protein 1 and specific component of the SET1A/B complex, serves as global regulation on H3K4 methylation. Thus, the elevation of CFP1 (Set1/COMPASS) suggests that KSHV infection increases the total level of H3K4me3 on the PROX1 promoter, and the recruitment of MLL2 (MLL2/COMPASS) resolves the bivalent domain of the PROX1 gene. Besides, no change in UTX (MLL3/4 COMPASS) was detected in the PROX1 gene after KSHV infection (Fig. 4D). RBBP5 and WDR5, shared by all six methyltransferase complexes (Set1A, Set1B, and MLL1-4), did not respond significantly to KSHV infection with a slight increase of RBBP5 (Fig. 4E) and a minor decrease of WDR5 (Fig. 4F). Therefore, we conclude that KSHV infection results in the recruitment of MLL2 and Set1 complexes to the PROX1 promoter region to elevate active histone marker H3K4me3.

**Fig. 4.**
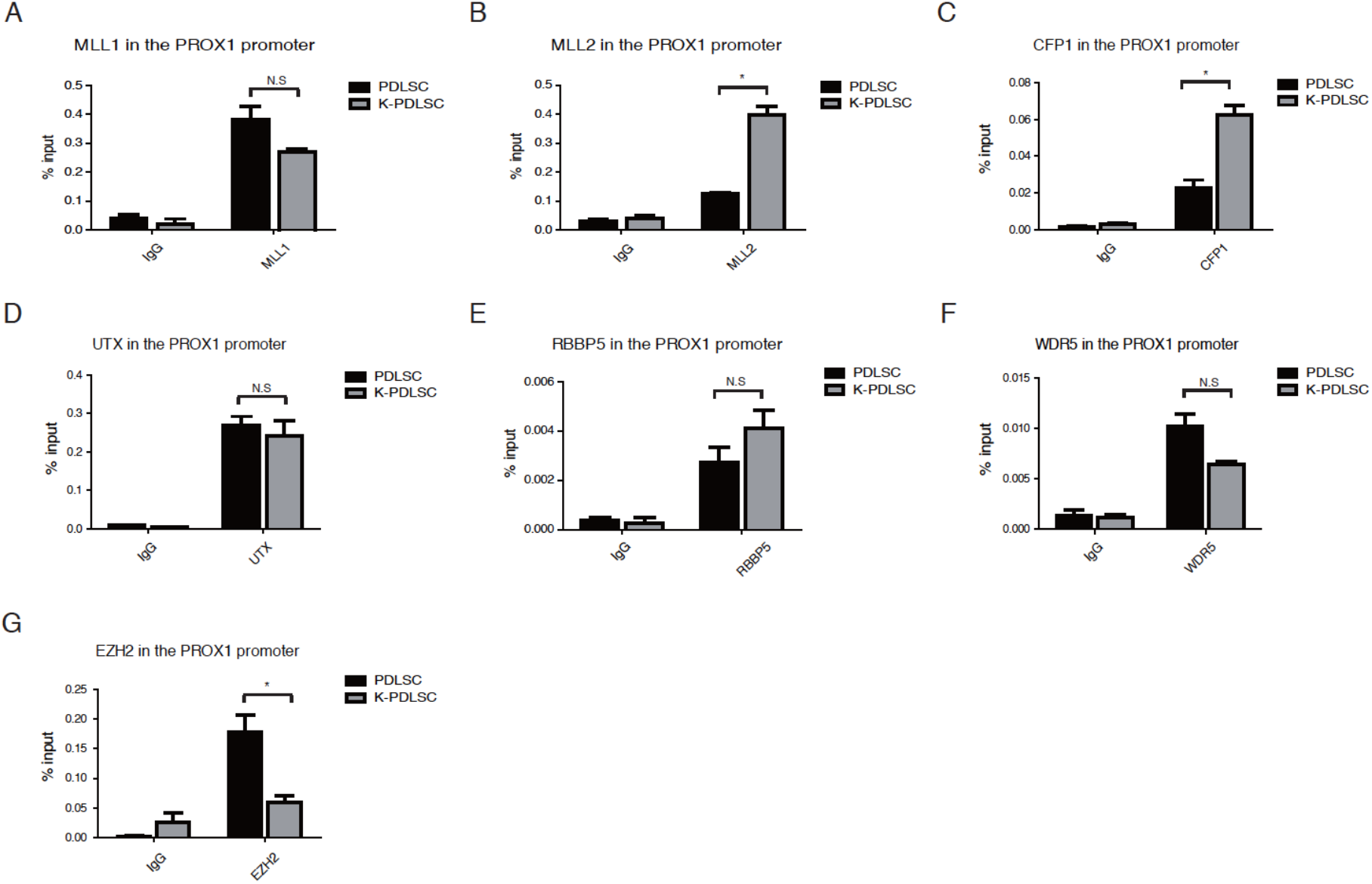
KSHV resolves the PROX1 bivalent chromatin by recruiting MLL2 and CFP1 and removing PRC2 to the PROX1 Promoter. Mock- and KSHV-infected PDLSCs were subjected to ChIP assay with antibodies against different histone methyltransferase components, including MLL1, MLL2, CFP1, UTX, RBBP5, WDR5, and EZH2 (a component of PRC2). Error bars represent SD. Statistics analysis was performed using the one-way ANOVA test. P-values < 0.05 were considered significant (*P<0.05, **P<0.01 and ***P<0.001) and N.S represents no significance.

On the other hand, the histone methyltransferase responsible for the deposition of the H3K27me3 mark on a bivalent domain is the PRC2 complex (Cao et al., 2002; Czermin et al., 2002; Kuzmichev et al., 2002). Our result showed that KSHV infection led to the removal of EZH2, the catalytic subunit of the PRC2 complex, from the PROX1 promoter, causing the loss of the H3K27me3 mark in the PROX1 region (Fig. 4G). Altogether, KSHV-mediated recruitment of MLL2 and Set1 and removal of PRC2 resolve the bivalent domain of the PROX1 promoter.

### Identification of KSHV components that are responsible for epigenetic regulation of PROX1 gene expression

To elucidate the mechanism underlying KSHV altering the epigenetic features of the PROX1 gene, we needed to identify the viral genes that control PROX1 activation. Inspection of KSHV transcription profiles of KS lesions (Rose et al., 2018) and KSHV-infected PDLSCs (Li et al., 2018) revealed a class of KSHV genes expressed in both KS lesions as well as KSHV-infected MSCs. We ectopically expressed these KSHV genes, including latent genes LANA, K12, and vFLIP and lytic genes K1, ORF45, ORF50, K8, PAN, vIL-6, and vGPCR in PDLSCs. Among these viral genes, vIL-6 and vGPCR significantly promoted PROX1 expression (Fig. 5A).

**Fig. 5.**
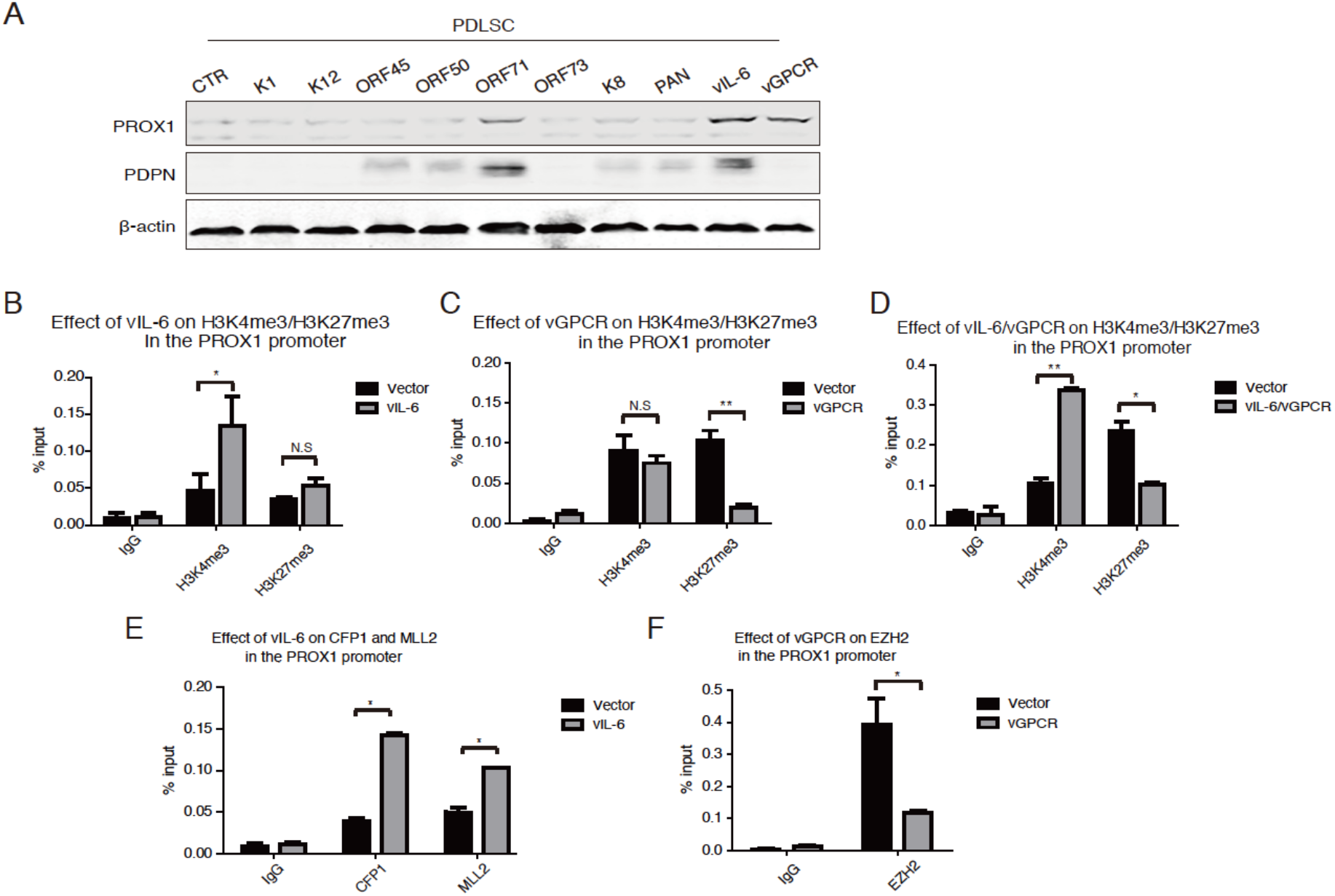
Identification of KSHV components that are responsible for the epigenetic regulation of PROX1 gene expression. (A) PDLSCs were transfected with a class of viral gene expression vectors, including K1, K12, ORF45, ORF50, ORF71, ORF73, K8, PAN, vIL-6, and vGPCR. The effects of these viral components on PROX1 expression were assayed by Western blot. ChIP assays were performed with PDLSCs ectopically expressing vIL-6 (B), vGPCR (C), or both (D) for their effects on H3K4 and H3K27 trimethylation in the PROX1 promoter. The effects of expression of vIL-6 (E) and vGPCR (F) on CFP1, MLL2, and EZH2 in the PROX1 gene were examined by ChIP assays. Error bars represent SD. Statistics analysis was performed using the one-way ANOVA test. P-value < 0.05 was considered significant (*P<0.05, **P<0.01 and ***P<0.001) and N.S represented no significance.

The KSHV-encoded homolog to human interleukin-6 (vIL-6) could transmit its signals through gp130 to initiate proinflammatory and neoangigenesis and be sufficient to induce LEC-specific markers—PROX1 and PDPN (Vart et al., 2007; Morris et al., 2008; Chen et al., 2009). Viral G-protein coupled receptor (vGPCR) could upregulate PROX1 during LEC reprogramming and render cells enhanced cell proliferation, migration, and tumor formation (Aguilar et al., 2012). To explore if vIL-6 and vGPCR promote PROX1 expression at the epigenetic level, we examined the effect of vIL-6 and vGPCR expression on histone modifications in the PROX1 gene in PDLSCs. We found that vIL-6 expression led to a significant increase in H3K4me3 marks but had no effect on H3K27me3 content, while vGPCR expression caused a dramatic decrease in H3K27me3 marks in the PROX1 promoter but did not change the level of H3K4me3 (Fig. 5B and C). Co-expression of vIL-6 and vGPCR reproduced the alterations of H3K4me3 and H3K27me3 observed in KSHV-infected PDLSCs and complete resolution of the bivalent domain (Fig. 5D). Furthermore, vIL-6 expression deposited H3K4me3 via recruiting MLL2 histone methyltransferases to the promoter, and vGPCR removed H3K27me3 from the bivalent domain by downregulating PRC2 complex (EZH2) in the PROX1 promoter (Fig. 5E and F). Taken together, KSHV resolves the PROX1 bivalent domain through two pathways, namely vIL-6 and vGPCR signaling, to alter active H3K4me3 and repressive H3K27me3, respectively, and consequently regulate PROX1 gene expression and MEndT process.

### KSHV-mediated vIL-6 and VEGF secretions regulate MSC differentiation by resolving the bivalent chromatin structure of the PROX1 gene

KS is an angiogenesis tumor, and pathological neoangiogenesis is a hallmark of the cancer. We previously found that the treatment of MSCs with the conditioned medium of KSHV-infected MSCs sufficiently renders cells increased angiogenesis activity and MEndT, indicating that KSHV can induce MSC MEndT in a paracrine manner (Zhong et al., 2017; Li et al., 2018). We examined the conditioned medium of KSHV-infected PDLSCs for its effect on PROX1 expression and found that the treatment of PDLSCs with the conditioned medium dramatically upregulated PROX1 expression in PDLSCs, especially 48 hours post-infection (48 hpi) (Fig. 6A). Furthermore, the treatment with conditioned media effectively altered the histone modification pattern exactly resembling what was observed in KSHV-infected PDLSCs, i.e., increased H3K4me3 and decreased H3K27me3 in the PROX1 promoter region (Fig. 6B).

**Fig. 6.**
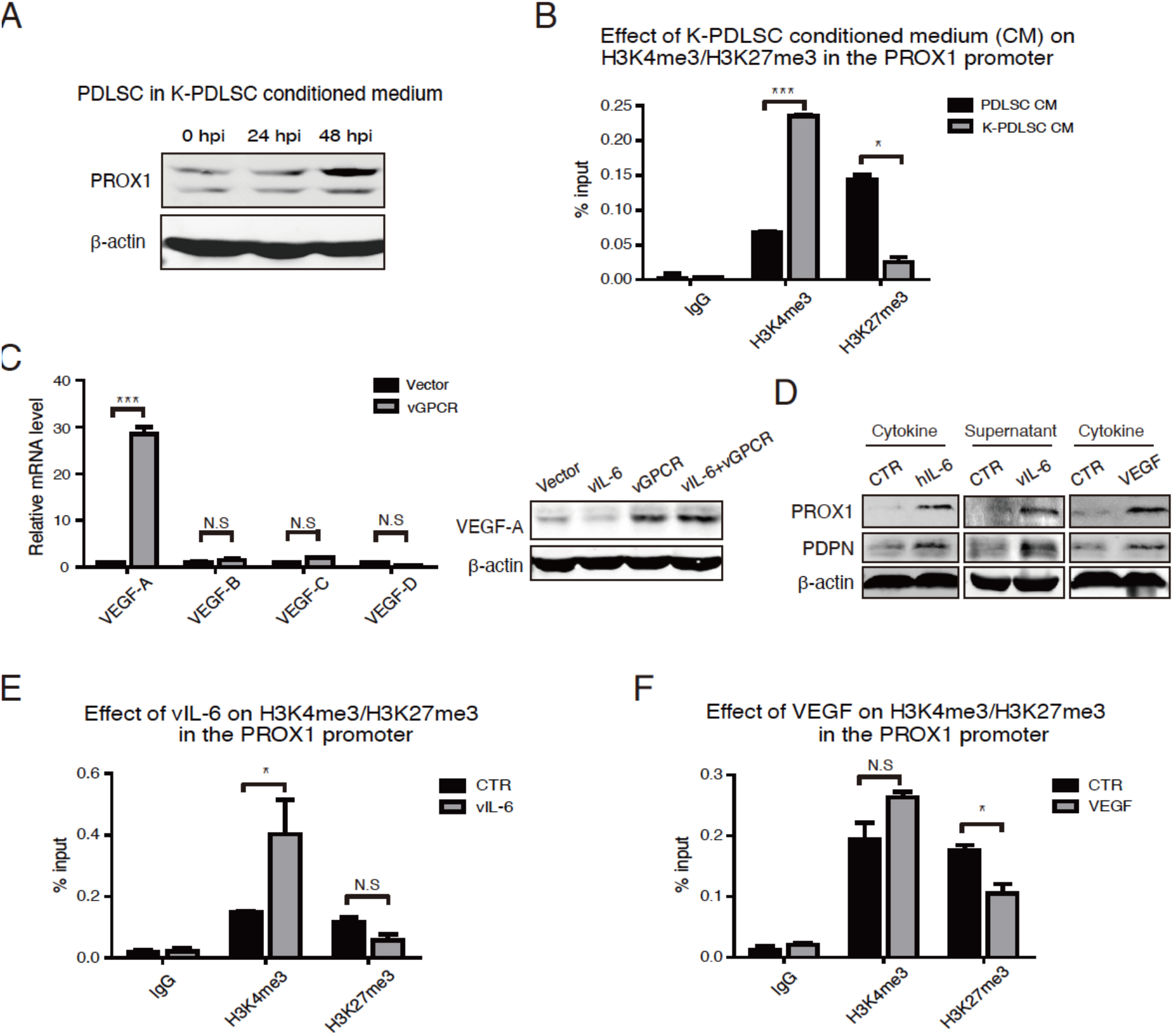
Cytokines mediate the epigenetic regulation of PROX1 expression in KSHV-infected PDLSCs. (A) PDLSCs treated with conditioned media of KSHV-infected PDLSC at indicated time points were examined for PROX1 expression by Western blot. (B) The conditioned medium-treated cells were subjected to a ChIP assay for histone modifications (H3K4me3 and H3K27me3) in the PROX1 promoter. (C) vGPCR was found to induce the production of VEGF-A, revealed at mRNA and protein levels. (D) Human interleukin-6 (hIL-6), viral interleukin-6 (vIL-6) supernatant of HEK293T cells transfected with a vIL-6 expression vector, and VEGF were added in PDLSC cultures, and cells were analyzed by Western blot with PROX1 antibody. ChIP assays were performed with vIL-6- (E) and VEGF-treated PDLSCs (F) for H3K4me3 and H3K27me3 modifications in the PROX1 promoter. The error bars represent SD. Statistics analysis was performed using t-test in GraphPad Prism. P-values < 0.05 were considered significant (*P<0.05, **P<0.01 and ***P<0.001) and N.S represented no significance.

In light of that KSHV-encoded vIL-6 and vGPCR can resolve the bivalent chromatin domain of the PROX1 gene, and so is the conditioned medium of KSHV-infected MSCs, we proposed a model to explain a possible mechanism for the paracrine regulation of epigenetic property of PROX1 gene as follows. It has been reported that vGPCR could upregulate VEGF production to induce angiogenesis and lymphatic reprogramming (Bais et al., 1998; Aguilar et al., 2012). Thus, vGPCR-induced VEGF, together with vIL-6, may be necessary and sufficient to resolve the bivalent chromatin and activate the PROX1 gene. We found that ectopic expression of vGPCR indeed induced VEGF-A expression in PDLSCs, as revealed by RT-qPCR and Western blot analysis (Fig. 6C). When PDLSCs were treated by human interleukin-6 (hIL-6), vIL-6 supernatant, and VEGF-A, we observed the up-regulation of PROX1 expression and its downstream PDPN expression (Fig. 6D). Finally, the effects of vIL-6 and VEGF-A on histone modification of the PROX1 gene were analyzed by treating PDLSCs with the supernatant of vIL-6-transfected 293T cells (48 hours post-transfection) and VEGF-A, followed by ChIP assay for H3K4me3 and H3K27me3. Interestingly, the treatment with vIL-6 supernatant led to increased H3K4me3, and VEGF-A caused decreased H3K27me3 in the PROX1 promoter (Fig. 6E and F). Therefore, we conclude that KSHV induces PROX1 activation through a dual signaling process, i.e., activating vIL-6 signaling to increase active histone marker H3K4me3 and using vGPCR-VEGF-A axis to reduce repressive histone marker H3K27me3, to resolve the bivalent chromatin structure and activate PROX1 gene expression.

### vIL-6 signaling and vGPCR-VEGF axis are indispensable for epigenetic regulation of PROX1 and acquisition of tumorigenic properties

To assess the significance of vIL-6 and vGPCR-VEGF axis-mediated epigenetic regulation for PROX1, we silenced these two viral genes individually or simultaneously in KSHV-infected PDLSCs by using small interference RNA (siRNA). Knockdown of vIL-6 and vGPCR, each moderately inhibited PROX1 expression compared to control siRNA (si-NC) (Fig. 7A and C). The inhibition of vIL-6 reduced H3K4me3 marks in the PROX1 promoter but did not affect the H3K27me3 marker (Fig. 7B), while knockdown of vGPCR expression increased H3K27me3 in the promoter but had no effect on H3K4me3 (Fig. 7D). Double knockdown of vIL-6 and vGPCR dramatically reduced PROX1 expression (Fig. 7E) by decreasing active mark H3K4me3 and elevating repressive mark H3K27me3 in the PROX1 promoter, counteracting the effect of KSHV infection in the PROX1 promoter (Fig. 7F).

**Fig. 7.**
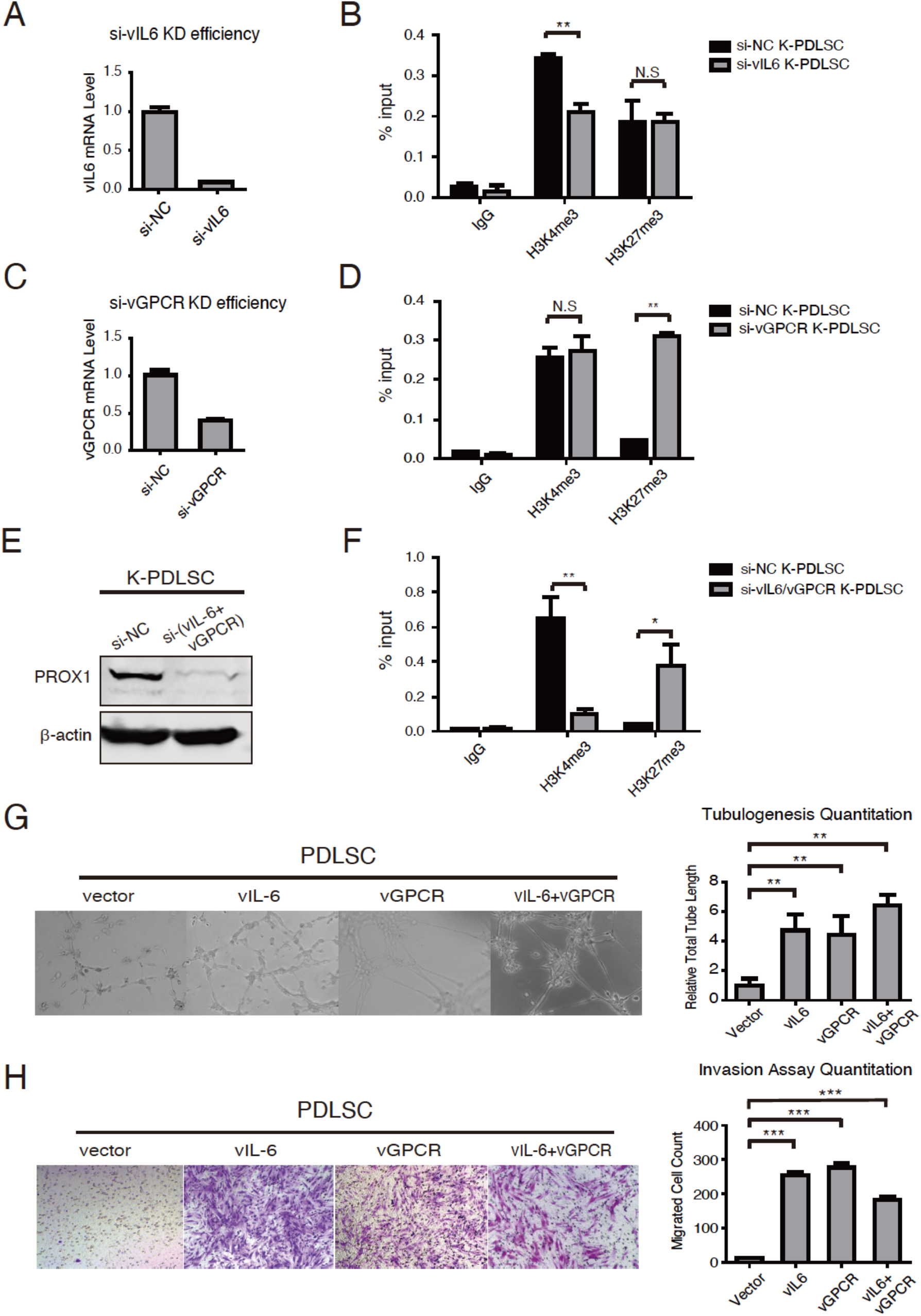
vIL-6 signaling and vGPCR-VEGF axis contribute to KS tumorigenic properties. (A) vIL-6 expression was silenced with siRNA (si-vIL-6 and negative control si-NC) in KSHV-infected PDLSCs. The knockdown efficiency was verified with quantitative RT-PCR. (B) KSHV-PDLSCs transduced with si-vIL-6 or si-NC were subjected to ChIP assays to evaluate H3K4me3 and H3K27me3 in the PROX1 promoter. (C) vGPCR expression was silenced with a specific siRNA (si-vGPCR) in KSHV-infected PDLSCs. The knockdown efficiency was verified with quantitative RT-PCR. (D) KSHV-PDLSCs transduced with si-vGPCR or si-NC were subjected to ChIP to evaluate H3K4me3 and H3K27me3 in the PROX1 promoter. (E) KSHV-infected PDLSCs co-transduced with si-vIL-6 and si-vGPCR were assayed for PROX1 expression by Western blot, and (F) the histone modifications in the PROX1 promoter by ChIP assay. (G) vIL-6, vGPCR, and vIL-6/vGPCR ectopically expressed PDLSCs were placed on Matrigel, and tubulogenesis was examined by Tubule formation assay and quantified by measuring the total segment tube length using the ImageJ software. (H) vIL-6, vGPCR, and vIL-6/vGPCR ectopically expressed PDLSCs (1.5×10^4^ cells/well) were seeded in the upper chamber of Transwell in serum-free α-MEM. α-MEM containing 20% FBS was used as a stimulus for chemotaxis in the lower chamber. Cells migrated to the lower chamber were stained with crystal violet. Statistics analysis was performed using the one-way ANOVA test, and P-value was calculated by GraphPad Prism. P-values < 0.05 were considered significant (*P<0.05, **P<0.01 and ***P<0.001) and N.S represented no significance.

Then the contribution of vIL-6 signaling and vGPCR-VEGF axis to the MEndT process and tumorigenic properties were evaluated. vIL-6 and vGPCR were reported to possess the potentials to promote tumorigenesis and angiogenesis (Bais et al., 1998; Aoki et al., 1999). In vIL-6 or/and vGPCR-overexpressed PDLSCs, angiogenesis activity represented by theformation of capillary-like tubules was found significantly increased (Fig. 7G). Using a Matrigel-transwell assay, we evaluated the invasive feature of vIL-6 or/and vGPCR-overexpressed PDLSCs. The result showed that vIL-6 and vGPCR expression increased the migration and invasion ability of PDLSCs (Fig. 7H).

## Discussion

We previously reported that KSHV infection of human MSCs triggers an MEndT process that may lead to Kaposi’s sarcoma development (Li et al., 2018). The current study sought to elucidate the mechanism of KSHV-mediated MEndT. The central event in viral-initiated MEndT is KSHV-mediated gene expression reprogramming that re-routes the differentiation process of MSCs. In this study, we found that the expression of the PROX1 gene, a master regulator of endothelial lineage differentiation and MEndT, is controlled at the epigenetic level through a bivalent chromatin domain that is usually seen in embryonic development transcription factor genes (Bernstein et al., 2006). KSHV infection activates PROX1 by resolving the bivalent domain through a dual signaling process, in which vIL-6 signaling increases active histone marker H3K4me3, and vGPCR-VEGF axis decreases repressive histone marker H3K27me3 (schematically depicted in Fig. 8). As a consequence, the activation of PROX1 initiates endothelial lineage differentiation and MEndT, leading to MSC acquisition of tumorigenic features such as proliferation, angiogenesis, and migration/invasion.

**Fig. 8.**
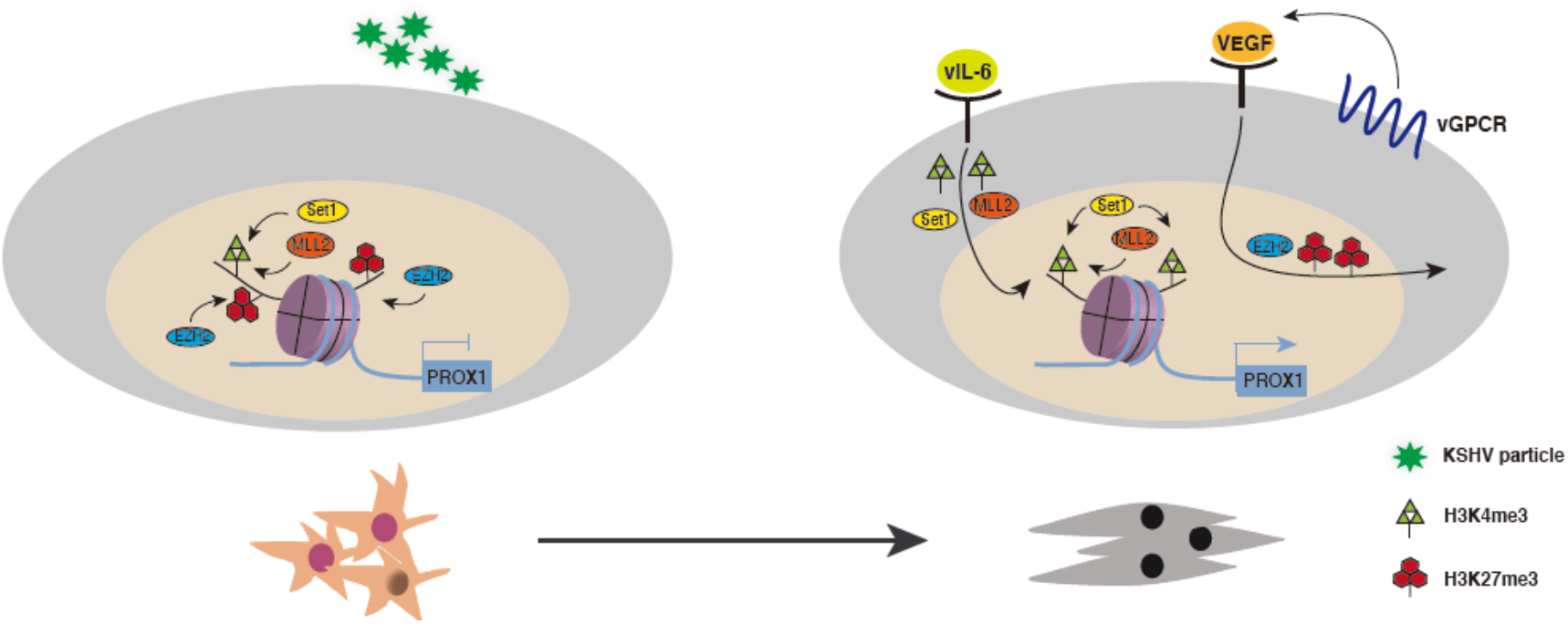
Model for KSHV-mediated epigenetic regulation of MEndT through the bivalent chromatin of PROX1 gene. PROX1 gene in MSCs harbors a distinctive “bivalent” epigenetic signature consisting of both active marker H3K4me3 and repressive marker H3K27me3. KSHV infection resolves the bivalent chromatin by decreased H3K27me3 and increased H3K4me3 to activate the PROX1 gene. vIL-6 signaling leads to the recruitment of MLL2 and Set1 complexes to the PROX1 promoter to increase H3K4me3, and the vGPCR-VEGF-A axis is responsible for removing PRC2 from the promoter to reduce H3K27me3. Therefore, through a dual signaling process, KSHV activates PROX1 gene and initiates MEndT, thereby transforming MSCs into KS-like spindle-shaped cells.

Postnatal stem cells have multi-lineage differentiation potential into the adipocytes, chondrocytes, or osteocytes (Pittenger et al., 1999). PROX1 is a homeodomain transcription factor essential for the development of a variety of organs, including the lymphatic system, lens, heart, liver, pancreas, and central nervous system (Oliver et al., 1993; Wigle et al., 1999). During early lymphatic development, endothelial cells in the cardinal vein exhibit a mixed phenotype of BECs and LECs. A subset of venous endothelial cells begins to express PROX1 and migrates out to form initial lymphatic vessels (Wigle et al., 2002). Overexpression of PROX1 in blood vascular endothelial cells (BEC) was able to induce lymphatic vascular endothelial cells (LEC)-specific gene transcription in the BECs (Petrova et al., 2002). KSHV infection of BECs and LECs reprograms both types of cells to move away from their original cell identities and acquire the opposite cell type (LEC and BEC, respectively) properties, in accompanying with up- and down-regulation of PROX1 expression, respectively(Hong et al., 2004; Wang et al., 2004). Further study delineated the opposing regulation of PROX1 by KSHV to direct cell differential fate through IL-3R (up-regulation) and Notch (down-regulation) signaling (Yoo et al., 2012). In our previous study, we found that KSHV-infection of oral MSCs can up-regulate PROX1 and, as a consequence, promote endothelial-lineage differentiation or MEndT, while KSHV-infection of terminally differentiated LECs down-regulates PROX1 (Li et al., 2018). Taken together, PROX1 is a key regulator in KSHV-mediated MEndT that reprograms MSCs leading to KS sarcomagenesis. In the current study, we confirmed the critical roles of PROX1 in developing KS tumorigenic phenotypes (Fig. 2). Therefore, the regulation of PROX1 expression by KSHV is a key event in transforming a KSHV-infected MSC to Kaposi’s sarcoma. The revelation of epigenetic regulation of PROX1 in MSCs by KSHV in this study provided a novel insight into the MEndT process and KS tumorigenesis.

Herein, the PROX1 gene was found to reside in bivalent chromatin where activating histone H3 Lys4 trimethylation (H3K4me3) mark and the repressive histone H3 Lys27 trimethylation (H3K27me3) mark co-exist on the same nucleosome. In 2006, a subset of promoters associated with both activating H3K4me3 and repressive H3K27me3 marks, referred to as “bivalent” chromatin domain, was discovered in mouse embryonic stem cells (ESCs) (Bernstein et al., 2006). Such a signature bivalent chromatin appears to be more specific to the promoter of developmentally important transcription factors. The synchronous existence of H3K4me3 and H3K27me3 posits these transcription factor genes in a poised state, enabling them to be rapidly activated upon suitable developmental cues and environmental stimuli (Voigt et al., 2013). Our finding that the transcription factor PROX1 gene is also posited in the distinctive bivalent chromatin in oral MSCs suggests that lineage differentiation transcription factor genes in postnatal stem cells are regulated at the epigenetic level in the same mechanism as that used in embryonic stem cells. In this study, we investigated how KSHV resolved the bivalent chromatin domain for PROX1 activation and revealed that KSHV uses a dual signaling process, i.e., activating vIL-6 signaling to increase active histone marker H3K4me3 and using vGPCR-VEGF-A axis to reduce repressive histone marker H3K27me3, to resolve the bivalent chromatin structure and activate PROX1 gene expression. This is an interesting finding and suggests that there is a “two-factor-authentication” mechanism in the resolution of bivalent chromatin. The biological significance for such a two-factor-authentication has not yet been understood, but we speculate that genes in bivalent domain chromatin are poised for a quick response to an external stimulus. Still the dual signaling process may grant a more efficient and accurate (with an additional layer of security) response to activate the gene.

vIL-6 and vGPCR were found to play essential roles in resolving the bivalent chromatin domain and activating the PROX1 gene. vGPCR exerts this function through producing VEGF-A. Thus, vIL-6 and VEGF-A regulate PROX1 expression through an autocrine or paracrine mechanism. This finding explained our previous observations that MSCs can acquire KS tumorigenic properties by being incubated with conditioned media of KSHV-infected MSCs (Zhong et al., 2017), and MSCs could undergo MEndT via paracrine regulation(Li et al., 2018). This finding also implies that KSHV may not only regulate PROX1 expression in its infected MSCs, but also modulate uninfected cells in the environment for PROX1 activation. In fact, it is well known that Kaposi sarcoma secrets abundant inflammation cytokines and growth factors including IL-1, IL-6, basic fibroblast growth factor (bFGF), platelet-derived growth factor (PDGF), tumor necrosis factor (TNF), interferon-gamma (IFN-γ), and vascular endothelial growth factor (VEGF), as well as KSHV-encoded vIL-6 (Boshoff et al., 1995; Jussila et al., 1998; Ensoli et al., 2001; Wang et al., 2004). In the early stage, the KS lesion is not a real sarcoma but an angiohyperplastic-inflammatory lesion mediated by inflammatory cytokines and angiogenic factors. With the development of KS, spindle-shaped tumor cells accumulate and become dominant, triggered or amplified by KSHV infection (Ensoli et al., 2000). Our result is consistent with the notion that KS is a cytokine disease and further suggests that cytokine autocrine or paracrine plays a central role in establishing KS spindle-shaped tumor cells in KS lesions. However, it is worthy of noting that although KSHV-triggered PROX1 activation initiates the MEndT process, other KSHV-encoded oncogenes and virally induced signaling may also contribute to MEndT and transformation of MSCs to KS cells at different stages.

## Materials and Methods

### Ethics statements

The use of human samples and PDLSCs in this study was approved by the Medical Ethics Review Board of Sun Yat-sen University (approval no. 2015-028). The animal experiments were approved by the Animal Ethics Review Board of Sun Yat-sen University (approval no. SYSU-IACUC-2018-000162) and carried out strictly following the Guidance suggestion of caring laboratory animals, published by the Ministry of Science and Technology of the People’s Republic of China.

### Cell culture

The isolation of human PDLSCs has been described previously (Li et al., 2018). PDLSCs were maintained in alpha minimal essential medium (α-MEM) with 10% heat-inactivated fetal bovine serum (FBS), 2 mM L-glutamine and antibiotics. The second to tenth passages of PDLSCs were used in this study. Human embryonic kidney HEK293T cells were purchased from American Type Culture Collection (ATCC) and cultured in Dulbecco’s modified Eagle’s medium (DMEM) supplemented with 10% FBS and antibiotics. iSLK.219 cells, cultured in DMEM with 10% FBS and antibiotics, were gifted from Dr. Ke Lan at Wuhan University.

### Reagents and antibodies

Cell culture medium (α-MEM and DMEM) and FBS were purchased from GIBCO Life Technologies. Penicillin and streptomycin were obtained from HyClone. Glutamine was obtained from Sigma. Matrigel for tube assay and cell invasion assay was obtained from Corning. Transwell inserts (PIXP01250) were purchased from Merck Millipore. Anti-PROX1 (BA2390) for Western blot and anti-PROX1 (ab199359) for immunohistochemistry and immunofluorescence were purchased from Boster and Abcam, respectively. Anti-LANA (ab4103) antibody for immunohistochemistry and immunofluorescence was purchased from Abcam. Antibodies against H3K4me3 (07-473) and H3K27me3 (07-449) were from Merck Millipore, and methyltransferases CFP1 (CXXC1,40672), EZH2 (5246), MLL2 (63735), WDR5 (13105) and RBBP5 (13171) were from Cell Signaling Technology, Inc. Antibody against UTX (ab36938) was from Abcam. Antibody against MLL1 (61295) was purchased from Active Motif. Human IL-6 (hIL-6) (14-8069-62) was from eBioscience, and VEGF-165 (100-20) was obtained from PeproTech, Inc. Antibody against VEGF-A (YT5108) was from ImmunoWay. PDPN (11629-1-AP), CD31 (11265-1-AP), VEGFR2 (26415-1-AP), VCAM1 (11444-1-AP) were from ProteinTech. LYVE-1 (sc-19316) and VEGFR3 (a5605) were obtained from Santa Cruz and Abclonal, respectively.

### KSHV preparation and infection

iSLK.219 cells carrying rKSHV.219 were induced for lytic replication by 1 μg/mL doxycycline and 1 mM sodium butyrate. Five days post-induction. The culture media were filtered through a 0.45 um filter and ultra-centrifuged with Beckman Coulter OptimaTM L-100XP at 100000 g for 1 h. The pellet was resuspended in 1/100 volume (culture media) of 1x PBS and stored at −80°C until use. PDLSCs were seeded at 2×10^5^ cells per well in 6-well plates. Cells were infected with KSHV in the presence of polybrene (4 μg/mL) at an MOI of 50 (viral genome copy equivalent). After centrifugation at 2500 rpm for 60 min at room temperature, the cells were incubated at 37°C with 5% CO_2_ for 2 hours. Then, the inoculum was removed by changing culture medium.

### Chromatin immunoprecipitation (ChIP)

PDLSCs (1×10^7^ cells) were cross-linked with formaldehyde of 1% final concentration for 15 min at room temperature, and the reaction was stopped by adding 125 mM glycine. Cells were washed with cold 1x PBS three times and collected by scratching the cells with cold 1x PBS. The cells were spun down under 2000 g at 4°C, resuspended with 1 mL lysis buffer (1% SDS, 10 mM EDTA, 50 mM Tris-HCl PH7.0, freshly added 1 mM PMSF, complete protease inhibitor [Roche]) for 10 min at 4°C and subjected to sonication to fragment the genomic DNA in size range of 300-700 base pair. Samples were centrifuged at 14000 g for 15 min at 4°C, and supernatants were collected as soluble chromatin. For each ChIP assay, 100 μl of chromatin solution was diluted to 1 ml with dilution buffer (0.01% SDS, 1.1% Triton X-100, 1.2 mM EDTA, 20 mM Tris-HCl pH8.0, 167 mM NaCl, complete protease inhibitor [Roche]). Antibody (2 μg) was added to the chromatin sample and incubated overnight at 4°C. Streptavidin magnetic beads (Merck Millipore), which had been washed three times in lysis buffer and blocked with 100 mg/mL sheared salmon sperm DNA (Ambion), were added to chromatin-antibody mixture and incubated at 4°C for 4 hours. The beads were washed 3 times with low salt (0.01% SDS, 1% Triton X-100, 2 mM EDTA, 20 mM Tris-HCl PH8.0, 150 mM NaCl, complete protease inhibitor [Roche]), high salt (0.01% SDS, 1.1% Triton X-100, 1.2 mM EDTA, 20 mM Tris-HCl PH8.0, 500 mM NaCl, complete protease inhibitor [Roche]), and LiCl (250 mM LiCl, 1% NP-40, 1% sodium deoxycholate, 1 mM EDTA, 20 mM Tris-HCl PH8.0, complete protease inhibitor [Roche]) washing buffers, respectively, and two times with 1x TE buffer (100 mM Tris-HCl, pH8.0 and 10 mM EDTA, pH8.0). DNA was eluted from the beads with 400 μl elution buffer (1% SDS, 0.1 M NaHCO_3_). Immune complexes were eluted by 150 ml TE buffer (pH8.0) and cross-links were reversed by heating at 65°C overnight, followed by digestion with Proteinase K (0.2 U/mL) at 50°C for 2 h. DNA was purified with Magen HiPure DNA Clean Up Kit (D2141) and analyzed by real-time PCR with primers for the PROX1 promoter.

### Sequential Chromatin immunoprecipitation (sequential ChIP)

Sequential ChIP was performed as previously described (Bernstein et al., 2006). Briefly, the first step was standard ChIP with antibody against H3K27me3 and the sheared DNA fragments were controlled in the range of 400-500 base pairs. Eluted chromatin was diluted 50-fold and subjected to a second immunoprecipitation with antibody against H3K4me3. Immune complexes were eluted, cross-linking was reversed, and DNA was purified as described above.

### Quantitative PCR (qPCR) and reverse transcription- qPCR (RT-qPCR)

PCR primers for evaluating ChIP or sequential ChIP assays were designed to amplify 150-200 base pair fragments from the promoter regions as follows: PROX1 (forward: 5’-GACCCCCAGATTCCCAGGTCCTTCT-3’; reverse: 5’-AAGCCAGATTTCTATATTTTTTCTG-3’), PML (forward: 5’-TTTCGGACAGCTCAAGGGAC-3’; reverse: 5’-TTAGTTTCGATTCTCGGTTT-3’), TGFBR2 (forward: 5’-AGCTGTTGGCGAGGAGTTTC-3’; reverse: 5’-AGGAGTCCGGCTCCTGTCCC-3’), and TGFB3 (5’-GGGAGTCAGAGCCCAGCAAA; reverse: 5’-TGGCAACCCTGAGGACGAAG-3’). Real-time PCR was carried out using SYBR Green PCR mix (Roche) in Roche LightCycler 480II Instrument.

RNA was isolated using an Ultrapure RNA Kit (CWBIO CW0581), reverse transcribed (Takara), and quantified using SYBR green PCR master mix on a Roche LightCycler 480II. The following primers (5’-3’) were used: VEGF-A (forward: AGGGCAGAATCATCACGAAGT; reverse: AGGGTCTCGATTGGATGGCA), VEGF-B (forward: GAGATGTCCCTGGAAGAACACA; reverse: GAGTGGGATGGGTGATGTCAG), VEGF-C (forward: GAGGAGCAGTTACGGTCTGTG; reverse: TCCTTTCCTTAGCTGACACTTGT), VEGF-D (forward: ATGGACCAGTGAAGCGATCAT; reverse: GTTCCTCCAAACTAGAAGCAGC), vIL-6 (forward: TCGTTGATGGCTGGTAG; reverse: CACTGCTGGTATCTGGAA), and vGPCR (forward: AACCATCTTCTTAGATGATGAT; reverse: AATCCATTTCCAAGAACATTTA).

### Immunohistochemistry (IHC) analysis

Paraffin-embedded Kaposi sarcoma samples and kidney capsule transplants were sectioned. Deparaffinated sections were subjected to hematoxylin and eosin (H&E) staining (Servicebio G1002). For IHC staining, sections were subjected to antigen retrieval using 10 mM sodium citrate buffer (pH 6.0) for 10 min with an electric pressure cooker. Then tissues were treated in 3% hydrogen peroxide for 10 min to quench the endogenous peroxidase activity and probed with antibodies against anti-PROX1 (1:200), LANA (1:150), CD31 (1:100), PDPN (1:100), or VCAM1 (1:100) at 4°C overnight. The primary antibody binding was detected using a goat anti-rabbit HRP secondary antibody (Maxim DAB-1031), followed by colorimetric detection using metal enhanced DAB. Tissues were counterstained with hematoxylin.

### Immunofluorescence assay

Cells were fixed with 3.6% formaldehyde in PBS for 10 min, permeabilized in 0.1% Triton X-100 for 15 min, and blocked in 1% BSA for 1 h at room temperature. The samples were incubated with antibodies against LANA (1:200) or PROX1(1:200) respectively at 4°C overnight. Goat anti-Rat IgG Alexa Fluor 555 (1:200; Invitrogen A-21434) and Goat anti-Rabbit IgG Alexa Fluor 647 (1:200; Invitrogen A-21244) were used as secondary antibodies. Nuclei were stained by Hoechst 33258 (Sigma) for 3 min at room temperature. Slides were examined under a Nikon pE-300 fluorescent microscope (400x), and three channels were recorded sequentially.

### Western blot

Cells were lysed on ice for 30 min with lysis buffer (50 mM Tris-HCl pH 7.4, 150 mM NaCl, 1% NP-40, 1 mM NaF, 1 mM Na_3_VO_4_, 1 mM PMSF, protease inhibitor cocktail [1 tablet in 50 mL lysis buffer, Roche]). The whole cell extract was denatured, resolved by SDS-PAGE, and transferred onto nitrocellulose membranes. The membranes were blocked in 5% nonfat milk in 1xTris-buffered saline (TBS) for 1 h and incubated with diluted primary antibodies PROX1 (1:1000), PDPN (1:1000), VEGFR2 (1:500), VEGFR3 (1:500), LYVE-1 (1:100), or VEGF-A (1:1000) at 4°C overnight. β-Actin (1:5000; Sigma A5441) served as the internal reference. IRDye 680LT/800CW goat anti-rabbit or anti-mouse antibody (Li-Cor Biosciences) was used as a secondary antibody. The blots were visualized in an Odyssey system (Li-COR).

### shRNA-mediated knockdown of PROX1 expression

An shRNA lentiviral vector targeting the CDS region of PROX1 mRNA (Clone ID: NM_002763.3-531s21c1) was purchased from Sigma-Aldrich. Lentiviral particles were prepared by transfecting HEK293T cells with pLKO.1-shPROX1 (or pLKO.1-shControl), psPAX2, pMD2.G plasmids at the ratio of 5:3:2. Media containing lentiviruses were harvested at 48 h and 72 h and used to transduce PDLSCs. Transduced PDLSCs were selected under 2 μg/mL puromycin for a week.

### Tube formation assay

48-well plates were pre-coated with matrigel (1:1 dilute with α-MEM without FBS, 100 μL/well) and incubated at 37°C for 1 h to allow gelation to occur. 1×10^5^ PDLSCs were suspended in 200 μL α-MEM without FBS and placed to the top of the gel. The cells were incubated at 37°C with 5% CO_2_ for 6-8 h, and tube formation images were captured using a ZEISS microscope. The quantitation of the tube was carried out using the ImageJ software to measure the length of the total tube segments in the image. The average value was used for the histogram.

### Cell invasion assay

Cell invasion assays were carried out in 24-well Transwell units. Briefly, polycarbonate filters with 12-mm pores were precoated with 100 μL of matrigel (1:10 diluted with α-MEM without FBS). 1.5×10^4^ cells in serum-free media were placed in the top wells, and the bottom chambers were filled with 20% FBS medium. After 24 h incubation, the cells that had passed through the filter were stained with crystal violet. The number of invaded cells was counted from multiple randomly selected fields using ImageJ software. Photographs were taken and independent experiments were performed in triplicate.

### Ectopic expression of viral genes

KSHV open reading frames including K1, K12, ORF45, ORF50/RTA, ORF71/vFLIP, ORF73/LANA, K8, PAN, vIL-6, and vGPCR were cloned into the pMSCVpuro vector at the Bgl II and EcoRI sites using ClonExpression II One Step Cloning Kit (Vazyme C112-02). Retroviral particles were prepared by transfecting HEK293T cells with pMSCVpuro-(K1, K12, ORF45, ORF50/RTA, ORF71/vFLIP, ORF73/LANA, K8, PAN, vIL-6, and vGPCR) and PIK plasmids at the ratio of 1:1. Media containing retroviruses were harvested at 48 and 72 h and used to transduce PDLSCs. Transduced PDLSCs were selected under 2 μg/mL puromycin for a week.

### siRNA-mediated silence of vIL-6 and vGPCR

Small interference RNAs targeting KSHV encoded interleukin-6 (vIL-6) and G-protein coupled receptor(vGPCR) were obtained from Guangzhou RiboBio Co., Ltd. Si-vIL6 (5’-TGTTCTGAGTGCAATGGAA-3’) and si-vGPCR (5’-CCTCATAAATGTTCTTGGA-3’) (or si-NC, si-GAPDH, Cy3) were transiently transfected into KSHV-PDLSCs using Lipofectamine 3000 Transfection Regent (Invitrogen L3000015). The transfection efficiency of siRNA was assessed according to Cy3 under a fluorescence microscope, and transfected cells were ready for the next experiment 48-72 h post-transfection.

### Supernatant transfer assay

The supernatant from KSHV-PDLSCs (48 hpi) was transferred to PDLSC culture at the ratio of 1:1. HEK293T cells were transfected with vIL-6 expression vector, and media were collected after 48 h post-transfection 48. The supernatant with vIL-6 was added to PDLSC culture at the ratio of 1:1.

### Kidney capsule transplantation

PDLSCs were suspended with α-MEM media and seeded on 96-well plates precoated with 0.5% agarose at 2×10^4^ cells per well. The spheroids were grown at 37°C for 1 to 2 d with 5% humid CO_2_. The media containing spheroids were collected and transferred to 15 mL conical tubes, washed twice with 1x PBS, and centrifuged at 1000 rpm for 5 min. About 100 spheroids were placed on gelfoam scaffold for 1 d in medium. Recipient female nude mice (6-8 weeks, n=3-5) were weighed and anesthetized with isoflurane. Kidney was exposed through an incision of the skin and muscle on the back of the mouse. The kidney capsule was opened with the fine tip of no.5 forceps. The spheroids/scaffold block was placed under the kidney capsule. Sutures were placed, and capsule-grafting products were harvested 28 d after transplantation.

### Statistical analysis

Data were analyzed by two-tailed Student’s t-test and one-way ANOVA in GraphPad Prism. P values < 0.05 were considered significant (*P<0.05, **P<0.01 and ***P<0.001) and N.S represented no significance.

## Acknowledgments

We thank all members of Yuan Lab for critical reading of this manuscript and constructive suggestions. This work is supported by grants from the Natural Science Foundation of China ((81530069).

## Author contributions

Y.D. and Y.Y. designed the research. Y.D., W.C., and Z.L. performed the experiments. Y.D. and W.C. performed the kidney transplantation of animal models. Y.W. provided human sample tissues. Y.D. and Y.Y. analyzed the data. Y.D. wrote the draft. Y.Y. wrote and reviewed the manuscript.

